# Feeding recombinant Royalactin: Measures of improved lifespan and mitochondrial function in *Caenorhabditis elegans*

**DOI:** 10.1101/2020.09.14.296053

**Authors:** Xia Li, Thomas L. Ingram, Ying Wang, Kamila Derecka, Nathan Courtier, Mattia Piana, Ian C. W. Hardy, Lisa Chakrabarti, Reinhard Stöger

## Abstract

Ageing, the decline of biological functions over time, is inherent to eukaryotes. Female honeybees attain a long-lived queen phenotype upon continuous consumption of royal jelly, whereas restricted supply of this nutritional substance promotes the development of worker bees, which are short-lived. An abundant protein found within royal jelly is major royal jelly protein 1 (MRJP1), also known as ‘Royalactin’. Health- and lifespan promoting effects have been attributed to Royalactin in species from diverse animal taxa, suggesting it acts on phylogenetically conserved physiological processes. Here, we explore the effects of feeding the nematode *Caenorhabditis elegans* with *Escherichia coli* that express a recombinant form of Royalactin (RA^rec^). We confirm that consumption of RA^rec^ increases body size, improves locomotion and extends lifespan. We discover a link between Royalactin and mitochondria, organelles which play a key part in the ageing process: both spare respiratory capacity and morphology indicate improved mitochondrial function in RA^rec^ fed *C. elegans*. These results demonstrate the feasibility of using recombinant Royalactin to gain further insight into processes of healthy ageing in many species.

RA^rec^ production allows insight into potential beneficial effects across species.

## 1 Introduction

Multicellular animals experience a decline of physiological functions over time and ultimately die. Evolutionarily conserved pathways and molecular networks are affected by these ageing processes, with mitochondrial dysfunction considered to be a common denominator [1,2]. Organisms have characteristic average lifespans and the comparative biology of long- and short-lived organisms offers a route towards untangling biochemical variations which account for lifespan differences [3,4]. Remarkable interspecific differences in lifespan can also be observed. For example, two types of adult females with divergent average lifespans develop in honeybee (*Apis mellifera*) colonies [5]. Queens live on average for 2-3 years and have larger bodies than their short-lived worker bee sisters, which live for only 3-6 weeks during the summer months and a little longer during winter periods [6,7]. The queen-worker dimorphism is dependent on the amount of royal jelly consumed during larval development: only those female larvae that are continuously fed royal jelly develop into queens, and royal jelly remains the exclusive food source of adult queens throughout their lifes [5].

Royal jelly has emerged as a compound of interest in longevity research and has been shown to extend the health and lifespan in evolutionarily distant organisms, including mice [8–10], the fruit fly *Dropsophila melanogaster* [11–13], crickets and silkworms [14] and the nematode *Caenorhabditis elegans* [15,16]. It is a secretion from the hypopharyngeal and mandibular glands of nurse bees and consists of a mixture of proteins, lipids, sugars and vitamins [17,18]. Royalactin (RA), also known as Major royal jelly protein 1 (MRJP1) is one of the proteins present in royal jelly[19] and has been postulated to be the key determinant of dimorphism and longevity differences among female honey bees [12]. While these claims are controversial [20,21], RA has been shown to decrease developmental time, increase size and increase longevity of the fruit fly *D. melanogaster* [12]. Similarly, feeding RA extracted from royal jelly extends longevity and enhances movement of *C. elegans* [22]. RA appears to exert its effects, in part at least, through signalling of the Epidermal Growth Factor (EGFR) pathway, in both honeybees and *C. elegans* [12,22]. RA was reported to improve longevity by also maintaining protein translation and proteasome activity following upregulation of chaperonins and elongation factors [23]. More recently, RA was found to act as a pluripotency factor, conferring self-renewal and promotion of a pluripotent gene network in mouse embryonic stem cells (mESC) [24]. A mammalian structural analogue of RA, Regina, has been identified and this protein is sufficient to promote ground-state pluripotency in mESCs [24].

Detienne and colleagues previously extracted RA from royal jelly by multiple rounds of ultracentrifugation, followed by reversed-phase high-performance liquid chromatography (RP-HPLC) to study its effects in *C.elegans* [22]. Here, we have fed *C. elegans* with bacterially expressed, recombinant RA (RA^rec^). Here, we show that feeding *C. elegans* with bacterially expressed recombinant RA (RA^rec^) positively affects body size, locomotion and lifespan. For the first time we find that RA^rec^ fed *C. elegans* have improved mitochondrial function, suggesting a common mechanism for the analogous effects observed across species.

## 2 Materials and methods

### 2.1 Preparation of bacterial food source containing Royalactin

The BL21 (DE3) *E. coli* strain containing the Lemo21 system was transformed with a pET28b+RA vector for expression of His-tagged RA (Lemo21+pET28b+RA) [25]. Expression cultures were incubated at 37°C until OD_600_ reached 0.6. RA^rec^ expression was induced by addition of 1M IPTG and overnight incubation at 37°C. To determine successful RA^rec^ expression samples were taken 0, 1, 2, 3, 4, 5 and 24 hours after incubation and subsequently stained on an SDS-PAGE gel (NuPAGE, Novex) using a His-Tag In-Gel stain (InVision) (Supplementary Figure 1). Stained gels were imaged at 302nm using an ImageQuant 300. As a negative control, Lemo21 System *E. coli* carrying an empty pET28b vector (Lemo21+ pET28b), which has no known target in *C. elegans*, was included in all experimental work. *E. coli* (Lemo21+ pET28b+RA and Lemo21+ pET28b) were pipetted onto NGM plates and used within a week.

### 2.2 Preparation of NGM agar

*Caenorhabditis elegans were* were maintained on nematode growth medium (NGM) agar using standard techniques [26]. To induce RA^rec^ protein expression, IPTG was added to liquid NGM at a final concentration of 400µM. A negative control group excluding IPTG was included. The antibiotics chloramphenicol and kanamycin were added to liquid NGM at a final concentration of 30μg/ml to maintain selection for Lemo21. *C. elegans* were raised on either *E. coli* strain OP50, Lemo21+ pET28b+RA or Lemo21+pET28b agar plates from L1 stage and were transferred to new plates each day.

### 2.3 Age synchronisation

The *C. elegans* strain CB5600 was used for control in all experiments and were handled as recommended by Sulston and Brenner [27], they were age synchronised by gravity flotation [28]. Gravity flotation was performed in M9 buffer (22.04mM KH_2_PO_4_, 42.27mM Na_2_HPO_4_, 85.56mM NaCl, 1mM MgSO_4_). A single 9.5 cm plate with high numbers of L1 larvae was washed at least three times with M9 buffer and transferred to a 15ml tube. L1 larvae were moved to a new NGM plate and incubated overnight at 20°C.

### 2.4 Lifespan experiments

All lifespan experiments were maintained at 20°C on NGM plates supplemented with kanamycin and chloramphenicol. Age-synchronised L1 *C. elegans* were kept on 95 mm NGM plates until they reached L3-L4 stage. Twenty to thirty *C. elegans* were transferred to each assay plate (35 mm diameter), corresponding to a total of 100 individuals per group. During the initial lifespan experiment, *C. elegans* were transfered onto fresh NGM agar plates every 24 hours. Lifespan was monitored daily starting from the first day of the adult L4 stage (scored as day 0). *C. elegans* that died were scored and any lost individuals were treated as censors.

### 2.5 Movement assays

The swimming activities of *C. elegans* were measured at adult days 7 and 14 as described by Zdinak *et al* [29]. Briefly, a single adult *C. elegans* was picked into 50μl M9 buffer on a microscope slide. The number of left and rightward body bends in ten seconds were counted, where one leftward and one rightward bend were one complete movement (stroke). A mean average of body strokes was calculated from a total of five individual trials. For each group the swimming activity assay was replicated 14 times, with 30 worms per replicate.

### 2.6 Measurement of body size and imaging of mitochondrial networks

*C. elegans* were transferred into 20μl M9 buffer on a glass slide and a cover slip applied. Images were taken using a DCF350FX camera mounted on a Leica IL 5000B upright microscope with a 5×/0.12 objective lens. Body sizes were measured using the image-analysing software WinROOF (Mitani corporation). CB5600 strain *C. elegans* express GFP in their mitochondria and muscle wall nuclei. *C. elegans* mitochondrial networks were imaged using a DCF350FX camera mounted on a Leica IL 5000B upright microscope with a 100×/1.30 oil-immersion objective lens.

### 2.7 High-resolution respirometry (HRR)

*C. elegans* grown on RA+IPTG or RA-IPTG were subjected to high-resolution respirometry (HRR) performed in a two-chamber system O_2_k-Respirometer (Oroboros Instruments, AT). One hundred live *C. elegans* were added to 0.52ml small volume glass chambers containing mitochondrial respiration media (MiR05-Kit, Oroboros Instruments, AT), at 20°C with a stirring speed of 750rpm. The output signal from the oxygen sensors were recorded at sampling intervals of 2 seconds using DatLab software (Oroboros Instruments, AT). The respirometer was calibrated at air saturation, at 20°C, before commencing experiments. High resolution respirometry (HRR) was carried out with a substrate-uncoupler-inhibitor titration protocol (SUIT_O3_ce) as follows: digitonin (1mM), oligomycin (omy; 2.5μM), uncoupler CCCP (carbonyl cyanide m-chlorophenyl hydrazone; 0.5μM titration steps), rotenone (Rot; 0.5μM), antimycin A (Ama; 2.5μM). Residual oxygen consumption (ROX), evaluated after inhibition of CIII by Ama, was used for flux correction across all respiratory states.

O_2_ flux values were used to calculate respiration in different coupling control states, including ROUTINE, LEAK, maximal ET (electron transfer) capacity and Complex I inhibited maximal ET capacity (S-ETC). ROUTINE respiration refers to physiological respiration, LEAK refers to residual respiration after ATP synthesis inhibition, maximal ET capacity is non-physiologic maximal uncoupled respiration and is demonstrated experimentally by uncoupler titration (CCCP), Complex I inhibited maximal ET capacity is respiration due to oxidative side reactions remaining after inhibition.

ROUTINE respiration was measured after *C. elegans* were titrated into the chambers containing mitochondrial respiration media; Digitonin was used to permeabilise cells; Omy was used to measure LEAK respiration; Uncoupler CCCP was used to determine ET capacity, which is observed when an optimum uncoupler concentration for maximal flux is reached; Rot was used to inhibit Complex I for measurement of non NADH-linked respiration.

### 2.8 Statistical analyses

GraphPad Prism (8.1.2) was used to for statistical analyses. Kaplan-Meier survival curves were generated using lifespan data (Cox Proportional Hazard or log-rank tests, where appropriate; corresponding *p*-values are referred to as *p*_*cox*_ or *p*_*log-rank*_). Brood sizes were analysed using one-way ANOVA and Tukey’s HSD post-hoc test. Movement and body sizes were analysed using two-way ANOVA with multiple comparisons. For HRR, determination of the effects of ROUTINE, inhibitors, or uncoupler, were assessed using unpaired *t*-test. Values *p*<0.05 were considered significant.

## 3 Results

### 3.1 Recombinant RA significantly increases *C. elegans* lifespan

We expressed a recombinant form of RA (RA^rec^) from the vector pET28b+RA without further purification, in an *E. coli strain* (Lemo21) optimised for IPTG-induced expression of proteins. To exclude that a possible lifespan extension was not an effect induced by cryptic elements present in the plasmid, an empty pET28b vector negative control (Lemo21 pET28b -IPTG) was tested. *C. elegans* are normally fed on a different *E. coli* strain (OP50) in laboratory conditions, and therefore an untransformed OP50 was included in our experiment for reference. To exclude IPTG effects, an additional control carrying the pET28b+RA vector without IPTG was included (RA-IPTG). We observed that the mean lifespan of the -IPTG group was significantly decreased compared to the OP50 group (−8.20%; *p*<0.05) (Table 1). *C. elegans* fed recombinant Lemo21+pET28b+RA+IPTG (RA+IPTG) had an extended lifespan compared to OP50 and negative controls (Figure 1). RA+IPTG increased the mean lifespan by 25.71% and median lifespan by two days compared with the OP50 group (12 and 10 days, respectively) (Table 1).

**Table 1.** Survival assay statistical comparisons of *C. elegans* groups RA+IPTG, OP50 or negative controls. n describes the total number of tested worms. SEM represents standard error around the mean, ^*^*p*_*COX*_<0.05 ^****^ *p*_*COX*_<0.0001. ^a^Mean lifespan change compared to positive control group OP50.

| Condition | n | Mean lifespan<br>(days ± SEM) | Increase vs<br>control<sup>a</sup> | $p_{COX}$ | Median<br>lifespan<br>(days) |
| --- | --- | --- | --- | --- | --- |
| OP50 | 100 | 10.97 ± 0.386 | --- | --- | 10 |
| -IPTG | 100 | 10.07 ± 0.287* | -8.20% | 0.027 | 9 |
| RA-IPTG | 100 | 10.71 ± 0.392 | -2.37% | 0.4186 | 9 |
| +IPTG | 100 | 11.60 ± 0.406 | 5.74% | 0.1656 | 11 |
| RA+IPTG | 100 | 13.79 ± 0.5**** | 25.71% | <0.0001 | 12 |

**Figure 1.**
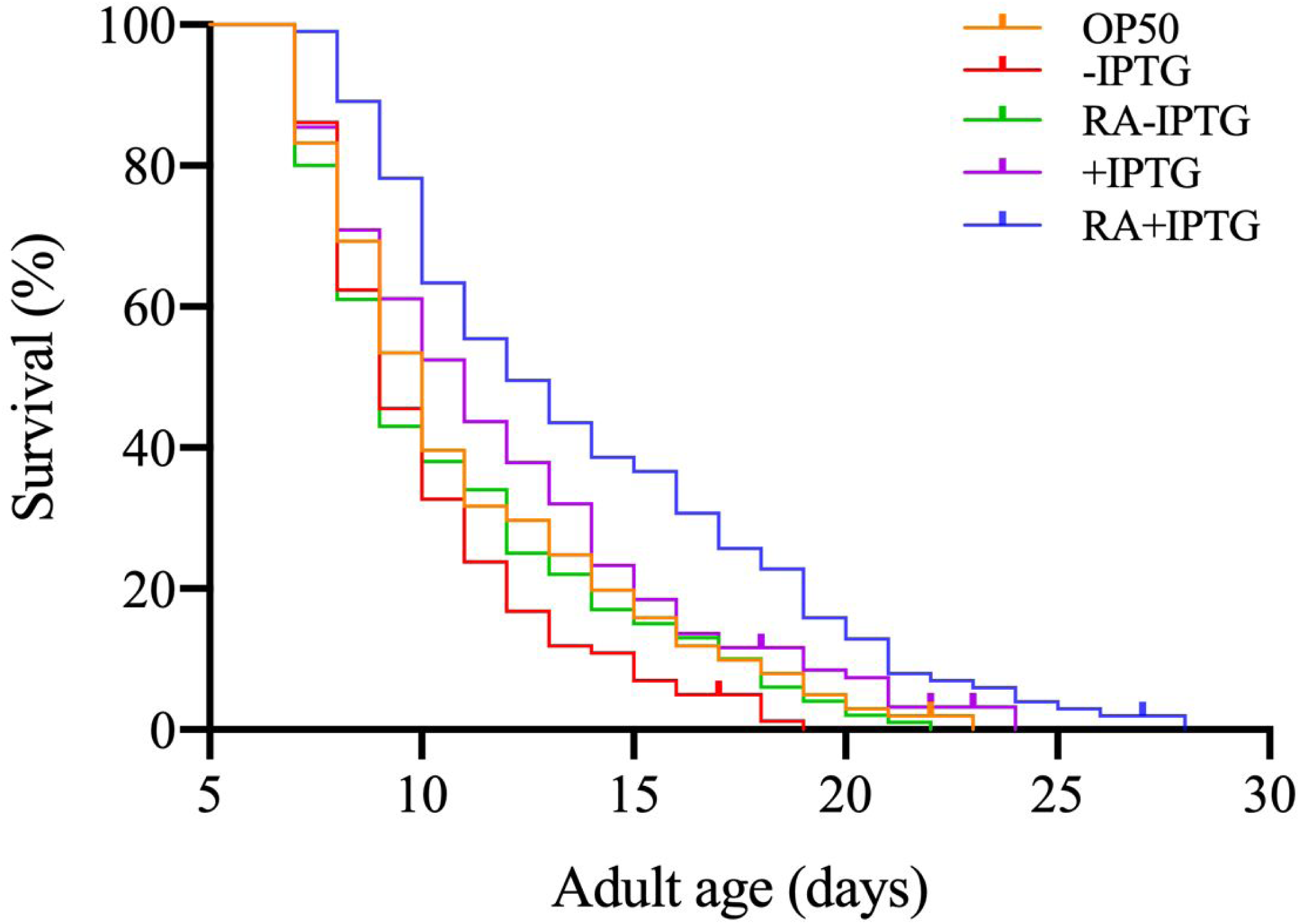
RA^rec^ fed worms have significantly increased mean and median lifespan compared to positive and negative control groups. The survival plot corresponds to parallel lifespan assays of the following treatment groups: positive control (OP50), negative control (-IPTG; +IPTG; RA-IPTG) and RA+IPTG *C. elegans*. IPTG was dissolved in NGM agar for +IPTG and RA+IPTG treatment groups. One hundred worms were used in each treatment group. Survival analyses were carried out by generating Kaplan-Meier survival curves (Cox Proportional Hazard or log-rank tests See Table 1 for statistical comparisons of treatment groups.

### 3.2 Royalactin increases the motility rate of *C. elegans*

We carried out a movement assay of 7- and 14-day old *C. elegans*, which were fed RA^rec^ expressing *E. coli* or controls. At day 7, RA+IPTG *C. elegans* had a significantly increased movement rate compared to negative control groups (*p*<0.05; 15.9% and 15.4% increase movement compared with -IPTG and RA-IPTG, respectively). No significant difference in movement rate was detected between RA+IPTG and OP50 *C. elegans*. Movement is known to decline with age in *C. elegans* [30] and we also observed a substantial age-related decrease in movement rate at day 14 compared with day 7 *C. elegans*. The age-related decrease was reduced in RA+IPTG *C. elegans* (Figure 2). RA+IPTG *C. elegans* had a significantly higher movement rate (26.5%) compared with OP50 group (*p*<0.05).

**Figure 2.**
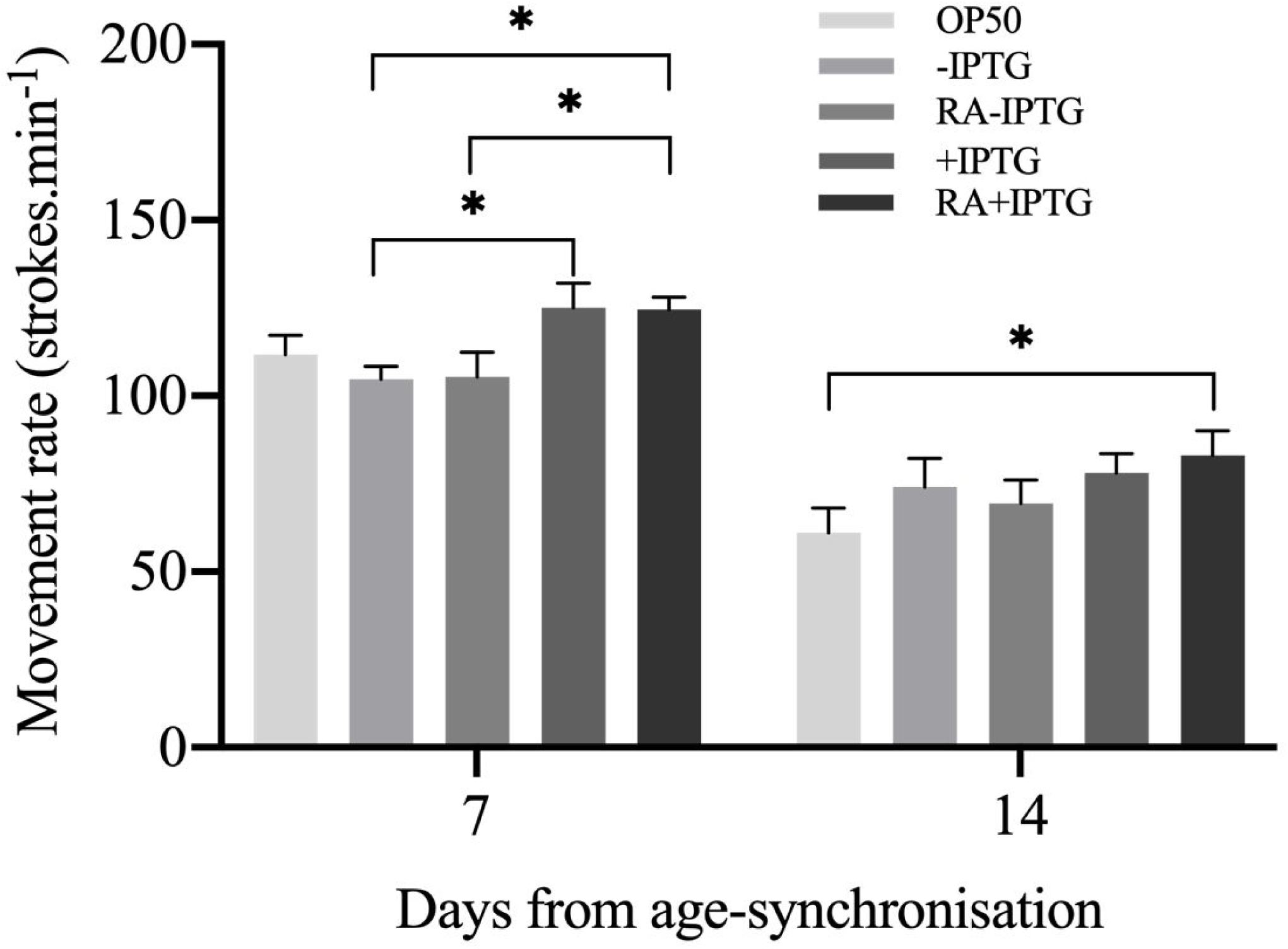
Effects of RA^rec^ on movement rate in *C. elegans* across the lifespan. Movement rates were measured at days 7 and 14 after age-synchronisation. RA+IPTG *C. elegans* have increased movement rate at day 7 compared to negative control groups (-IPTG and RA-IPTG). At day 14 RA+IPTG *C. elegans* show increased movement rate compared to OP50 group. Error bars represent SEM (two-way ANOVA with multiple comparisons), ^*^*p*<0.05. Each group consists of fourteen replicates with thirty *C. elegans* in each replicate.

### 3.3 Body size of *C. elegans* is increased with RA^rec^

We measured the effect of RA+IPTG on the body length of *C. elegans* at days 7 and 14 of adulthood. Adult *C. elegans* with RA+IPTG had increased body size (Figure 3A). The RA+IPTG group had significantly increased body length at days 7 and 14, compared to OP50 group (28.9% and 14.4% increased size; *p*<0.0001 and *p<*0.05, for days 7 and 14, respectively) (Figure 3B). In addition, the magnitude of body length increase of the RA+IPTG group compared to OP50 group was significantly greater between day 7 and day 14; this suggests RA accelerates early adulthood growth (mean body length differences were 0.45mm and 0.25mm, at days 7 and 14, respectively) (Figure 3B).

**Figure 3.**
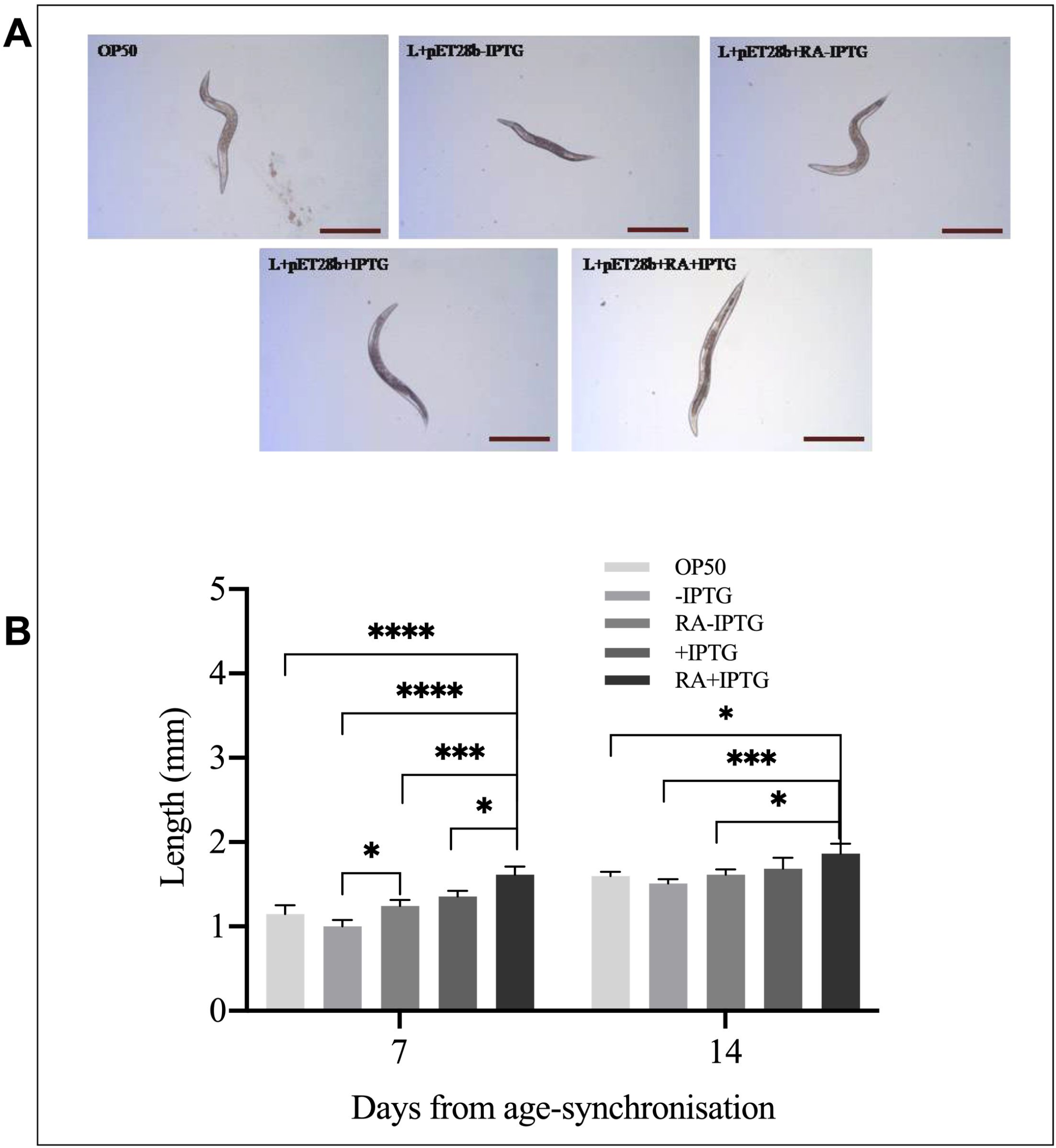
*C. elegans* fed RA^rec^ show significantly increased body size at days 7 and 14 compared to control groups. (A) Microphotographs of *CB5600 C. elegans* 8 days after hatching. Images were produced using a DCF350FX camera mounted on a Leica IL 5000B upright microscope with a 5×/0.12 objective lens. Scale bar is 500 μm. (B) Body length were analysed using image analysis software WinROOF (Mitani Corporation). Measurements of body length for each group were carried out twice (*n*=60 *C. elegans* per group). Error bars indicate SEM (two-way ANOVA with multiple comparisons). *\*p*<0.05 *** *p*<0.001 **** *p*<0.0001.

### 3.4 RA^rec^ fed *C. elegans* exhibit networked mitochondria

To assess mitochondrial morphological alterations due to RA^rec^ we imaged CB5600 muscle wall GFP-tagged mitochondria. CB5600 strain *C. elegans* contain mitochondria and nuclei that express GFP for visualisation. 8-day old worms from the control groups (OP50, -IPTG, RA-IPTG, +IPTG) have discrete mitochondria, with fragmented networks (Figures 4A-D). Mitochondria of 8-day old 8-day old RA+IPTG *C. elegans* are characterised by an elongated continuous morphology indicative of fused organelle networks (Figure 4E).

**Figure 4.**
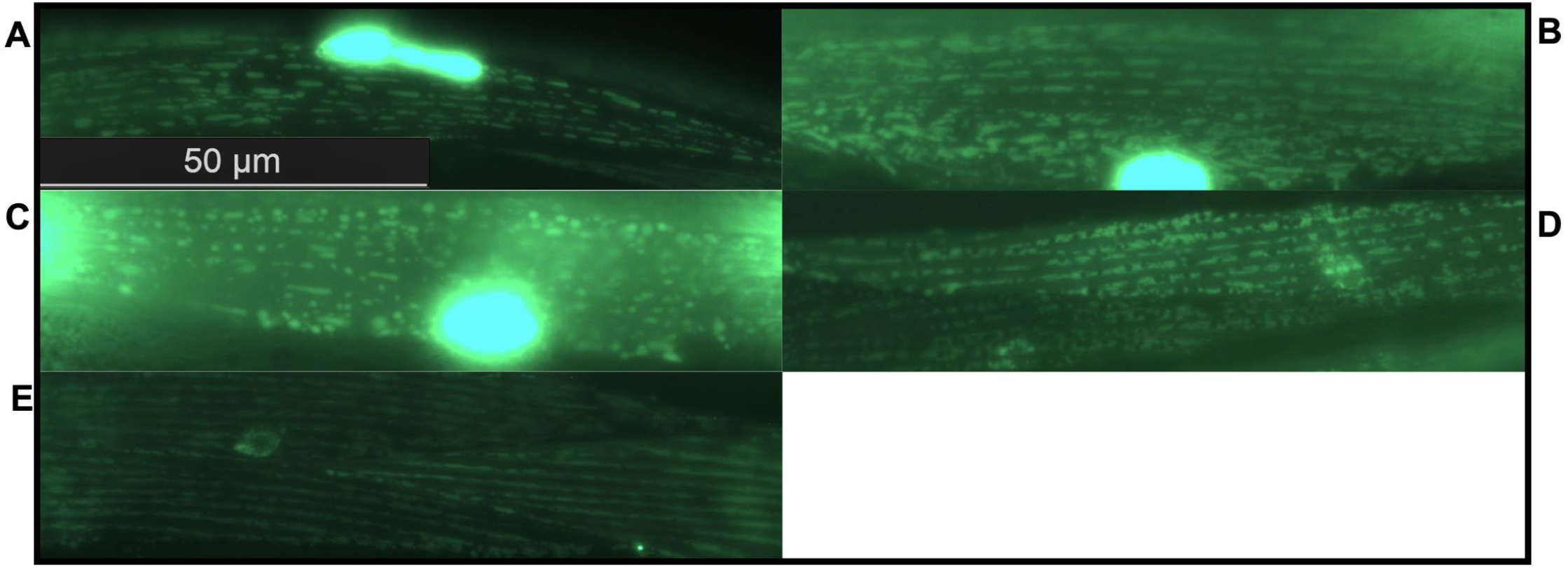
Mitochondria are elongated and orderly in RA^rec^ fed *C. elegans*. Fluorescence images of 8-day old *C. elegans* mitochondrial networks were produced using a DCF420 camera mounted on a Leica IL 5000B upright microscope, with a 100×/1.30 oil-immersion objective lens. A = OP50; B = -IPTG; C = RA-IPTG; D = +IPTG; E = RA+IPTG. The scale bar is 50μm.

### 3.5 RA^rec^ fed *C. elegans* have increased mitochondrial respiratory capacity

The mitochondrial respiratory function of RA^rec^ fed *C. elegans* was examined with substrate-uncoupler-inhibitor-titration (SUIT) protocols in an O2k Respirometer (Oroboros, AT) (Figure 5A). Experiments were run twice using RA+IPTG and negative control RA-IPTG groups only. Mean values for ROUTINE (R; *C. elegans* in MiR05-kit solution), LEAK (L; oligomycin addition), maximal electron transfer capacity (E; CCCP addition) and respiratory capacity through the S-pathway (S-ETC; rotenone addition) were not significantly different between groups. The mean respiratory magnitudes were, however, lower across RA+IPTG *C. elegans* (Figure 5B+C). Spare respiratory capacity, the difference between E and R respiration (E – R), describes the ability to produce additional ATP via oxidative phosphorylation during periods of increasing energy demand [31]. Spare respiratory capacity was significantly increased in RA+IPTG group compared with RA-IPTG (*p<*0.05).

**Figure 5.**
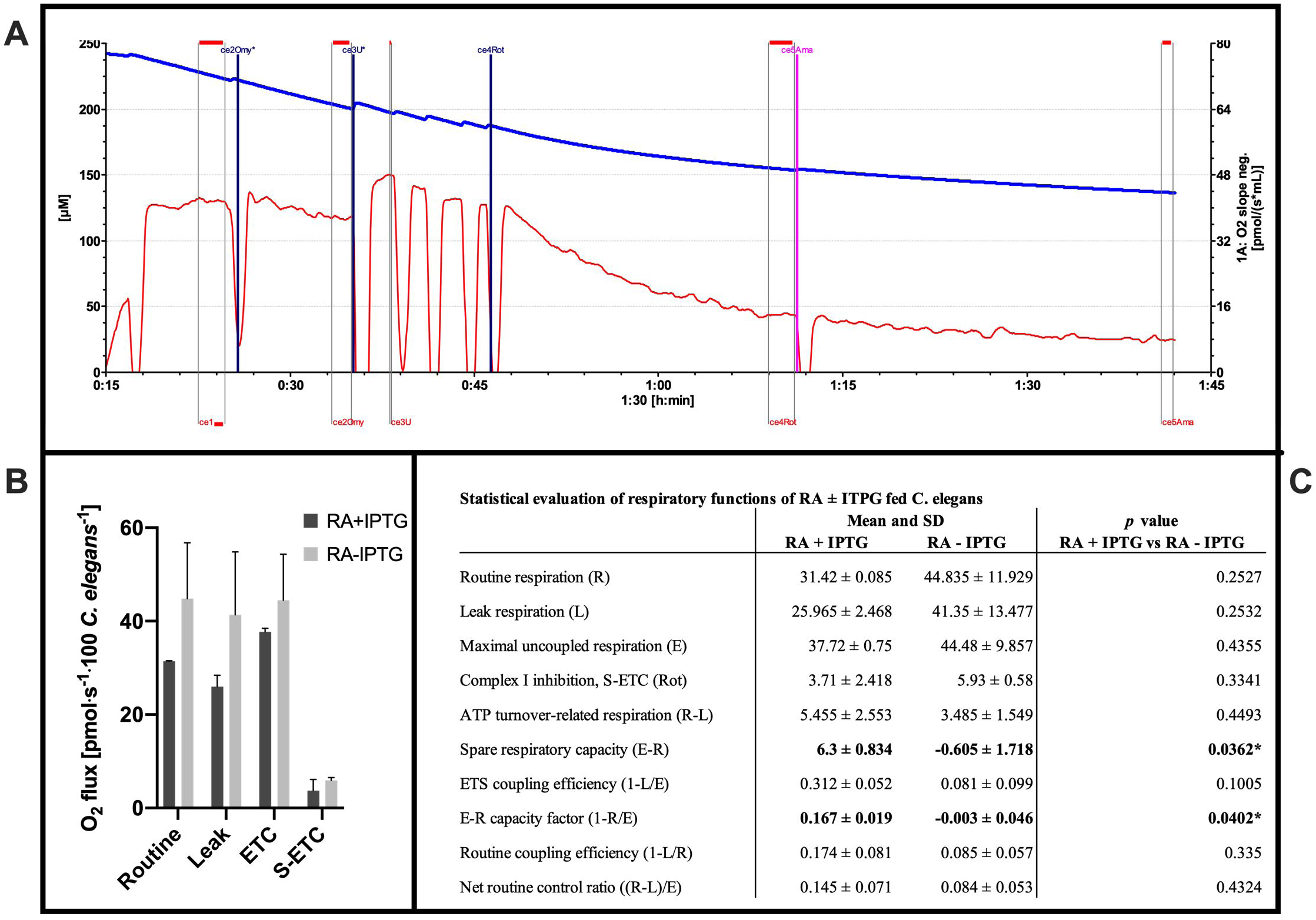
RA^rec^ increases reserve respiratory capacity of *C. elegans*. (A) Representative oxygraph trace of mitochondrial respiratory function of one hundred RA fed *C. elegans* in MiR05 respiration medium (Oroboros Instruments, AT) at 20°C, measured using SUIT_03_ce protocol (Datlab software). *C. elegans* were added to respirometer at the 15 minute mark. Omy = oligomycin; U = CCCP; Rot = rotenone; ama = antimycin A. Highlighted sections represent sampled oxygen flux regions. The red line represents oxygen consumption as oxygen flux per one hundred *C. elegans* (pmol O_2_ s^−1^ one hundred *C. elegans*^−1^); the blue line represents oxygen concentration (μM). (B) ROX-corrected respiration at 20°C. ROUTINE: one hundred *C. elegans*, no substrates or inhibitors; LEAK: oligomycin; ETC: electron transport capacity, stimulated by step-wise titration of CCCP; S-ETC: rotenone inhibition of Complex I, succinate-linked-ETC. Bar graphs represent mean ± SD (n = 2). (C) Mean ± SD and flux control factors ± SD of respiratory characteristics of RA-IPTG and RA+IPTG *C. elegans* groups (unpaired *t*-test) \**p<*0.05.

Flux control factors (FCF) were compared between RA+IPTG and RA-IPTG groups. FCF present a method for assessing the effect of an experimental variable on respiratory flux [32]. Excess E-R capacity factor, an expression of the relative scope for increasing R respiration in periods of increasing ATP demand, was significantly higher in RA+IPTG group than RA-IPTG (Figure 5C; *p<*0.05); E-R capacity factors were 0.167 ± 0.019 and -0.003 ± 0.046, corresponding to 16.7% and -0.3% of ETS capacity not being utilised during ROUTINE respiration, for RA+IPTG and RA-IPTG respectively. No significant differences in other FCF were observed (Figure 5C; mETS coupling efficiency, R coupling efficiency, net R control ratio).

## 4 Discussion

The average lifespan of humans has increased dramatically over the last 170 years [33] but longevity is often associated with the occurrence of debilitating medical conditions [34]. Progress in the field of ageing has revealed age-related biochemical changes that are conserved across animal species, and that modulating these changes impacts prevalence of age-related disease [35–38]. Several nutritional compounds, including Royal jelly, have been identified and appear to alter lifespan by modulating age-related pathways [12,22,39].

Royal jelly has been reported to increase the lifespan of species beyond the honeybees that produce it [5]. These include other insects, such as *D. melanogaster* [11–13] but also phylogenetically distant taxa such as mice [8,9,40] and *C. elegans* [15,16]. Isolated constituents of Royal jelly, including the lipid 10-HDA [16] and pantothenic acid [41], have also been shown to have effects. RA is reported sufficient to extend the lifespan and improve fertility of *D. melanogaster* [12], and prolong lifespan in *C. elegans* [22]. Our results using recombinant RA similarly find that bacterially expressed RA^rec^ is sufficient to extend the mean lifespan of *C. elegans* by over 25%.

A goal of ageing research is to match increased lifespan with healthspan, which can be defined as a period of life spent in good health [42]. Royal jelly, causes a faster growth rate of queens during development and increased body size of honeybee queens [6,7]. Here we have shown that feeding *C. elegans* recombinant RA significantly improved markers of healthspan. Feeding RA^rec^ increased the movement rate of *C. elegans* compared to controls at early-to-middle adulthood. Our results are in agreement with prior reports of improved movement following feeding on RA isolated from royal jelly [22]. In addition, feeding of RA^rec^ delayed the a*ge*-related decline in movement rate, suggesting RA^rec^ extends healthspan in *C. elegans*.

We have found that *C. elegans* fed RA^rec^ have increased body size in early-to-middle adulthood and that they are not larger simply because they reach maximal size more rapidly, but that they are larger throughout life. Although controversial, RA has also been reported to increase the rate of development and body size in the fruit fly *D. melanogaster* [12].

Mitochondrial dysfunction is a hallmark of ageing and age-related disease [1,43]. At day 8 of the worm lifecycle the mitochondrial network structure is generally fragmented and characteristic of age-associated muscle decline [44]. Interventions which promote longevity, such as exercise and caloric restriction, promote maintenance of mitochondrial function [45–47]. We have shown for the first time that RA^rec^ influences mitochondrial dynamics and function. Utilising the *C. elegans* strain CB5600, which contains GFP-tagged muscle wall mitochondria and nuclei, worms fed RA^rec^ have an elongated networked mitochondrial population. This phenotype has previously been associated with caloric restriction and was shown to protect mitochondria from autophagic degradation, thereby maintaining appropriate ATP levels for the cell under starvation conditions [48]. Chaudhari and Kipreos demonstrated that mitochondrial elongation is associated with numerous longevity pathways and that this phenotype is required for the survival of older animals during extended lifespans [49]. Additionally, high-resolution respirometry (HRR), we have demonstrated that RA^rec^ fed *C. elegans* have an increased spare respiratory capacity. Spare respiratory capacity, or respiratory reserve (RR), is associated with cellular stress response outcomes, where increased RR in cells improves resistance to cell death [50], while decreased RR is associated with ageing and increased risk of ageing-related diseases [51–53]. Together, our data suggest that RA^rec^ promotes mitochondrial functional states that are associated with longevity. We suggest that the further use of recombinant Royalactin will facilitate improvements in the understanding of how this protein promotes healthspan in diverse species.

## 5 Acknowledgments

This work was supported by the Biotechnology and Biological Sciences Research Council grant to Thomas L Ingram, a doctoral student [grant number BB/J014508/1]; China Scholarship Fund (CSC201806935060), China National Natural Science Foundation (11602159) supported Ying Wang and Xia Li. Mattia Piana’s PhD scholarship was jointly funded by Monkfield Nutrition Ltd. and the School of Biosciences / University of Nottingham.

## 6 Conflict of Interest

Monkfield Nutrition Ltd. sponsored in part the PhD scholarship of Mattia Piana. Otherwise the authors declare that the research was conducted in the absence of any commercial or financial relationships that could be construed as a potential conflict of interest.

## 7 Data Availability Statement

The original contributions presented in the study are included in the article/supplementary files, further inquiries can be directed to the corresponding author/s.

## Supplementary Figures

**Supplementary Figure 1.**
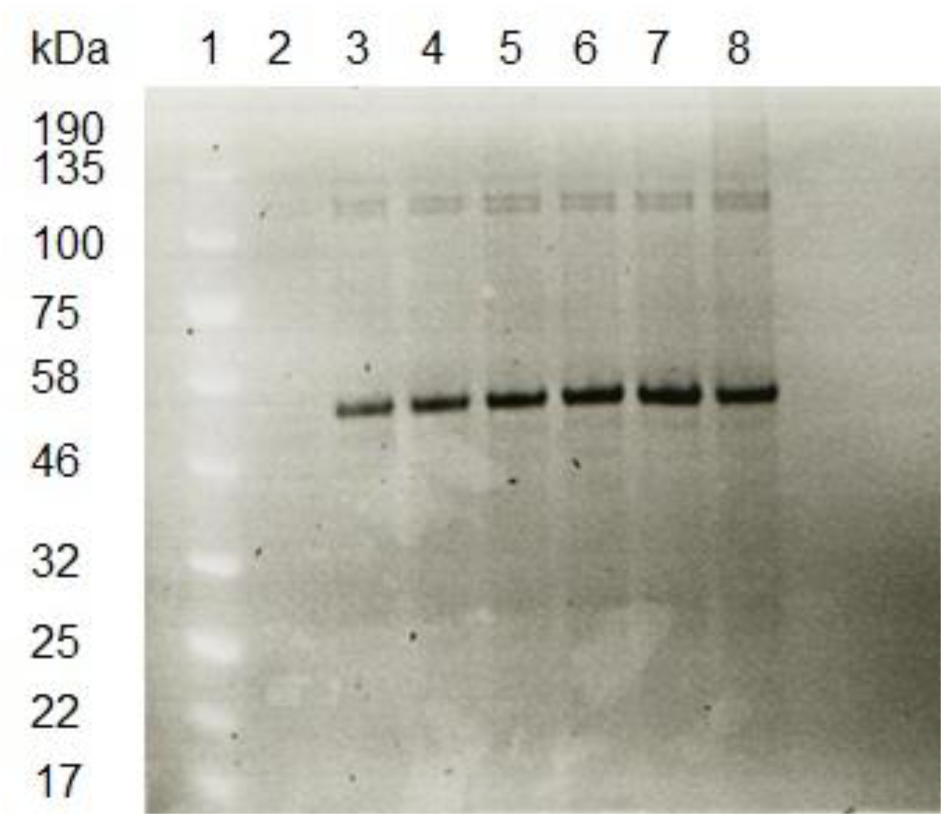
Culture samples induced to express His-tagged RA for 0 – 24 hours were run on a 4-12% Bis-Tris pre-cast SDS-PAGE gel (NuPage, Novex). Samples were prepared by adding 6 µL of sample to 3 µL of loading dye and 175 mM of DTT, followed by boiling at 90°C for 10 minutes. 10 µL of sample was loaded per well. Gels were run at 200 V for 1 hour 20 minutes, fixed (10% Acetic Acid, 40% Methanol) for 1 hour and stained with InVision His-Tag In-Gel Stain for 1 hour at room temperature. RA predicted molecular weight is 56kDa. The protein ladder used was Blue Protein Standards Broad Range (11 – 190 kDa) (NEB). Lanes: 1 = protein marker; 2 = 0 hour induction (T0); 3 = 1 hour induction (T1); 4 = 2 hour induction (T2); 5 = 3 hour induction (T3); 6 = 4 hour induction (T4); 7 = 5 hour induction (T5); 8 = 24 hour induction (T24). The Image was captured at 302 nm using an ImageQuant 300.

## Notes

### Competing Interest Statement

The authors have declared no competing interest.

